# AbiOmics: An End-to-End Pipeline to Train Machine Learning Models for Discrimination of Plant Abiotic Stresses Using Transcriptomic Profiling Data

**DOI:** 10.64898/2026.02.25.707868

**Authors:** Minkyu Park, Youngmin Oh, Woohyuk Choi, Yeong Deuk Jo

**Affiliations:** EuchroGene, LLC, Clarksburg, MD, 20871, United States; Department of Human-Centered Artificial Intelligence, Sangmyung University, Seoul, 03016, Republic of Korea; Department of Bio-AI Convergence, Chungnam National University, Daejeon, 34134, Republic of Korea; Department of Horticultural Science, Chungnam National University, Daejeon, 34134, Republic of Korea

**Keywords:** Abiotic stress, Transcriptomic profiling, Machine learning, Stress discrimination, *Arabidopsis*

## Abstract

Abiotic stresses are primary constraints on global crop productivity, reducing yields by up to 80%. While traditional phenotypic sensing detects stress only after physiological symptoms emerge and often fails to discriminate specific stressor types, transcriptomic profiling offers a high-dimensional solution, capturing rapid and sensitive molecular shifts. In this study, we developed AbiOmics, the first end-to-end machine learning pipeline specifically designed to identify and discriminate among multiple stressors. This approach represents a previously undocumented method for stress specification using large-scale transcriptomic big data. We identified 320 stress-specific marker genes using a curated collection of 1,243 transcriptomes of *Arabidopsis* samples treated with four major abiotic stresses, salt, cold, heat, and drought. A single-layer perceptron model trained on these features achieved 91% accuracy during five-fold cross-validation and 93% accuracy on an independent test set. The model demonstrated an unprecedented capacity to generalize to multi-stress conditions, identifying concurrent signatures in combinatorial salt-and-heat treatments. By integrating marker identification with SHAP-based biological interpretation, AbiOmics provides a rigorously validated diagnostic tool superior to conventional sensing. This framework establishes a high-confidence labeling strategy for AI-driven crop management and precision breeding to mitigate climate change impacts.

**Graphical Abstract:** 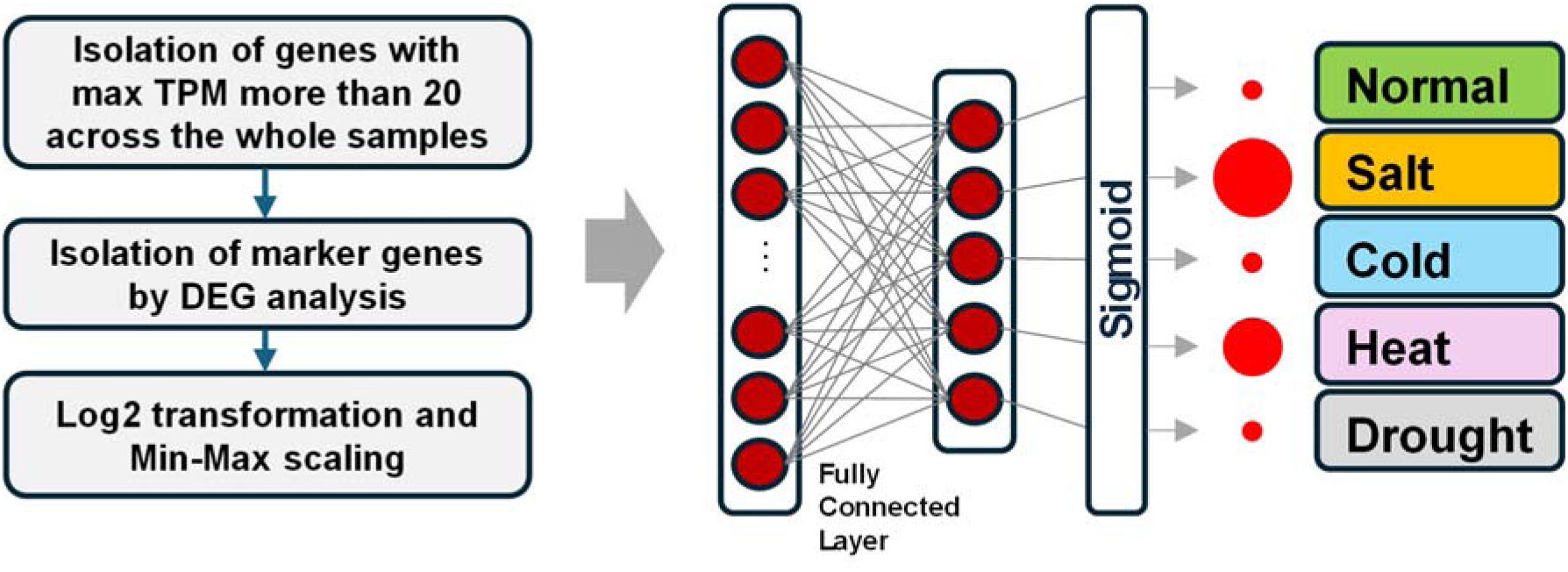

## Introduction

As sessile organisms, plants are inextricably linked to their environment, and adverse conditions during growth and development can cause severe tissue damage or mortality. Even under moderate suboptimal conditions, plants initiate sophisticated stress-signaling cascades that prioritize survival over productivity (1). This metabolic shift triggers fundamental changes in growth (2), organ development (3), senescence (4), and reproduction (5). For instance, plants employ reciprocal regulation through antagonistic interactions between intracellular regulators to balance stress responses with growth (6,7). This inherent trade-off significantly constrains biomass accumulation in unfavorable environments (8) , with research indicating that adverse conditions reduce average crop yields by 50%, and up to 80% in extreme cases (9–11).

Consequently, the precise identification of specific abiotic stressors is critical for elucidating adaptation mechanisms and developing tailored cultivation strategies to mitigate productivity losses.

Traditional stress diagnosis relying on visible phenotypic assessments is often inadequate, as discernible damage typically appears only after physiological decline is advanced. Moreover, distinct stressors often converge on similar phenotypes, complicating the identification of specific causal factors (12). To address these limitations, advanced imaging and sensing technologies have been deployed for early-stage diagnosis. Established methods leverage interpretable biological mechanisms, such as chlorophyll fluorescence imaging (CFI) for monitoring photosynthetic efficiency, thermal infrared (TIR) imaging for detecting stomatal closure under drought or heat, red-edge shift analysis for quantifying chlorophyll content, and LIDAR for evaluating structural architectural changes (13–17). While these techniques enable the detection of subtle physiological shifts, they are generally optimized for single stressors and lack the capacity to discriminate among multiple stressors simultaneously.

In contrast, hyperspectral imaging (HSI) captures reflectance across hundreds of narrow wavelength bands, facilitating the development of multi-stress diagnostic models through machine learning (18). However, while HSI is highly sensitive, its reproducibility is often compromised by environmental interference during measurement and the inherent complexity of data interpretation (12,18). Similarly, wearable electrochemical sensors have emerged as tools for real-time monitoring of tissue impedance or volatile organic compounds (VOCs) (19), yet they remain constrained by environmental noise and limited long-term stability. Despite these technological advancements, a significant bottleneck persists: most current methods can detect the presence of stress but fail to pinpoint the specific stressor type, making precise agricultural management difficult (20).

Transcriptomic profiling offers a promising solution to this challenge, as the transcriptome is intimately linked to stress-response mechanisms and undergoes rapid, sensitive shifts upon stress onset (12,21). Unlike phenotypic or spectral data, transcriptomes provide high-dimensional insights into the expression levels of every gene in the genome, potentially allowing for the discrimination of specific stressor types (22). While gene expression analysis has been used extensively to characterize stress-response pathways and identify tolerance factors (23,24), its application in stressor-specific diagnosis remains underdeveloped. Early efforts, such as a mini-scale microarray of 12 expressed sequence tags (ESTs) for identifying drought, salinity, and temperature stress, lacked rigorous validation and broad applicability (25).

Recently, the integration of machine learning with large-scale transcriptomic metadata has opened new avenues for stress diagnosis. Studies have analyzed hundreds of transcriptomes across various species, including *Arabidopsis* and barley, to identify core regulators of abiotic and biotic stress responses (26). Furthermore, machine learning models have successfully predicted disease severity in plant-pathogen interactions across diverse datasets (27). Despite these advancements, the use of transcriptomic big data to specify and distinguish between multiple abiotic stressors has not yet been reported.

In this study, we aimed to develop a robust machine learning model to identify specific abiotic stressors using transcriptomic metadata. To ensure broad applicability, we developed an end-to-end pipeline to train machine learning models for stress discrimination. We curated a comprehensive dataset of *Arabidopsis* leaf transcriptomes from public databases, covering key stressors: cold, heat, salt, and drought. Using these expression profiles, we trained and validated a diagnostic model. Subsequently, we evaluated its accuracy and generalizability using independent subsets of samples exposed to both single and combinatorial stress treatments.

## Materials and Methods

### Collection and processing of RNA-seq data

Transcriptomic datasets of *Arabidopsis* species subjected to abiotic stress were retrieved from the National Center for Biotechnology Information (NCBI) Sequence Read Archive (SRA).

Searches combined *Arabidopsis* with four abiotic stress terms: cold, heat, salt, and drought. Search results were filtered to include only datasets generated on Illumina sequencing platforms. To ensure data integrity, SRA accession numbers were manually curated by cross-referencing them with original published studies (see Results for detailed selection criteria). Raw SRA files were downloaded and converted to FASTQ format using the SRA toolkit. Because the retrieved datasets contained a mixture of single-end and paired-end libraries, all data were standardized to single-end format to ensure downstream compatibility; for paired-end libraries, only forward reads (R1) were retained.

Raw RNA-seq reads were quality-trimmed using AdapterRemoval (v2.3.4) with default parameters (28). Transcript quantification was performed using the *Arabidopsis thaliana* TAIR10 coding sequence (CDS) annotation as the reference (29). Read alignment and transcript abundance estimation (transcripts per million; TPM) were calculated using the RSEM pipeline (30) utilizing Bowtie2 for read mapping (31).

### Differential expression analysis, GO term enrichment, and marker gene selection

Differential expression analysis was performed on TPM values using the PyDESeq2 Python package. For each stress condition, 120 stress-treated samples were compared against 120 matched controls. Differentially expressed genes (DEGs) were identified using a threshold of |log_2_ fold change| (log_2_FC) ≥ 1 and an adjusted P-value ≤ 0.001. To minimize noise, transcripts with a maximum TPM < 20 across all samples were excluded.

Gene Ontology (GO) enrichment analysis was conducted using ShinyGO (v0.85.1) (32), with the Biological Process database. The genes for stress-specific up- and down-regulated DEGs were analyzed. Significant GO terms were identified using a False Discovery Rate (FDR) < 0.05. DEGs were categorized into up- and down-regulated groups, and Venn diagram analysis was employed to identify stress-specific DEGs for each of the four abiotic conditions.

To construct the diagnostic model, we established a consolidated set of 320 marker genes. This set was generated by randomly selecting 40 DEGs from each of the four up-regulated and four down-regulated stress-specific groups. Random selection was intentionally chosen over ranking-based approaches (e.g., by fold change or variance) to avoid overfitting to any single metric and to promote generalizability. The validity and robustness of this approach were subsequently confirmed by repeated sampling across 300 iterations (see below). The expression patterns of these 320 marker genes were visualized via heatmaps generated with the seaborn Python library.

To evaluate the robustness of marker selection, we assessed model performance stability across marker gene set sizes of 40, 80, 160, and 320. For each size, 300 random gene sets were generated, and performance was evaluated via 5-fold cross-validation and an independent test dataset. The variability in accuracy resulting from random selection is presented as distributions in violin plots. Furthermore, the concordance between cross-validation accuracy and independent test performance was analyzed using the Pearson correlation coefficient.

### Dimensionality Reduction Analyses

Principal component analysis (PCA) and t-distributed stochastic neighbor embedding (t-SNE) were performed using the scikit-learn Python library. Of the 27,416 genes detected across all samples, genes with TPM = 0 in all samples were removed, leaving 25,576 genes for dimensionality reduction analyses. For DEG-focused analyses, 6,670 non-redundant DEGs and the 320 selected marker genes were used. TPM values were log_2_-transformed and scaled using min–max normalization. A total of 1,243 curated RNA-seq samples, including 512 control, 148 cold, 133 salt, 266 heat, and 184 drought, were used for the analyses. Visualization of PCA and t-SNE results was performed using the seaborn Python library.

### Model training and evaluation

The model was trained using PyTorch. To prevent data leakage, the 65 independent test samples (13 per control group and 4 stress-treated groups) were fully excluded prior to all upstream processing steps, including DEG analysis and marker gene selection. The remaining 600 training samples (120 per control group and 4 stress-treated groups) were then split into a 5-fold cross-validation set using scikit-learn. Log_2_-transformed and min–max–scaled TPM values of the marker genes were used as model inputs. Training was performed with the following hyperparameters: Learning rate=0.005, Batch size=84, and optimizer=Nesterov-accelerated Adaptive Moment Estimation (NAdam). Optimal training epochs were determined using early stopping based on the minimum validation loss. Five models were trained, each corresponding to one cross-validation fold.

The importance of input marker genes was analyzed with SHapley Additive exPlanations (SHAP) (33). For each sample in the dataset, local SHAP values were calculated to quantify the impact of every marker gene on the individual prediction. The top 20 genes were ranked by their global importance scores, and their gene symbols and functions were retrieved using mygene 3.2.2, a Python package.

Model performance was evaluated using a 5-fold cross-validation test on the data and an independent test set of 65 samples. For single-stress samples, class predictions were based on the highest sigmoid output across the five classes (four stresses and one control). Performance was assessed using precision, recall, F1 score, and accuracy. Cross-validation performance metrics were averaged across folds, and standard deviations were calculated. For double-stress samples, sigmoid outputs for each stress class were visualized, with circle size representing the magnitude of the predicted probability.

## Results

### Collection and curation of stress-treated RNA-seq samples

To develop a machine learning model for discrimination of plant stress types, we curated RNA-seq datasets from *Arabidopsis* species subjected to four abiotic stresses: salt, cold, heat, and drought. An initial search identified 773 samples for salt stress, 640 for cold, 1103 for heat, and 942 for drought. Given the heterogeneity of experimental conditions of the samples, we applied stringent filtering criteria to ensure consistency and relevance for stress classification: 1) For drought stress, only samples induced by water deprivation were retained; those involving chemical inducers (e.g., salt) were excluded. 2) Samples involving additional treatments (e.g., hormones, pathogens, herbicides) were excluded. 3) Samples subjected to multiple simultaneous stresses were removed. 4) Only samples derived from leaf-related tissues (leaf, seedling, and rosette) were included. We did not filter based on stress duration, intensity, genotype, replication, or mutation background. Filtering was performed manually through a literature review. This process yielded 133 salt, 148 cold, 266 heat, and 184 drought stress samples. Control samples corresponding to each stress condition were also identified, totaling 82–183 samples.

To balance the dataset and prevent overfitting, we standardized the class counts. For each stress type, 120 samples were randomly selected for training and 13 for independent testing. From the pool of control samples, 30 per stress type (totaling 120) were selected for training, and 13 (3 each for salt, cold, and heat; 4 for drought) for testing. In total, 600 samples (120 per class) were used for five-fold cross-validation, and 65 samples were reserved for independent testing.

### Model training strategy

We employed a five-fold cross-validation strategy to ensure robust model training and evaluation (34). The dataset was split into training (80%), validation (10%), and test (10%) subsets. Early stopping was implemented based on validation performance (35). Test data was used to evaluate the trained models. As input data, normalized gene expression values (TPM) were used (Figure 1). Given the limited sample size (480 training samples), we performed differential expression analysis to reduce the input feature size. Models were trained on the selected genes and evaluated using cross-validation and independent test data. At the final stage, we evaluated the contribution of marker genes to model performance using SHAP. All these steps were constructed into an end-to-end pipeline for broad applications of the method to other plant species.

**Figure 1.**
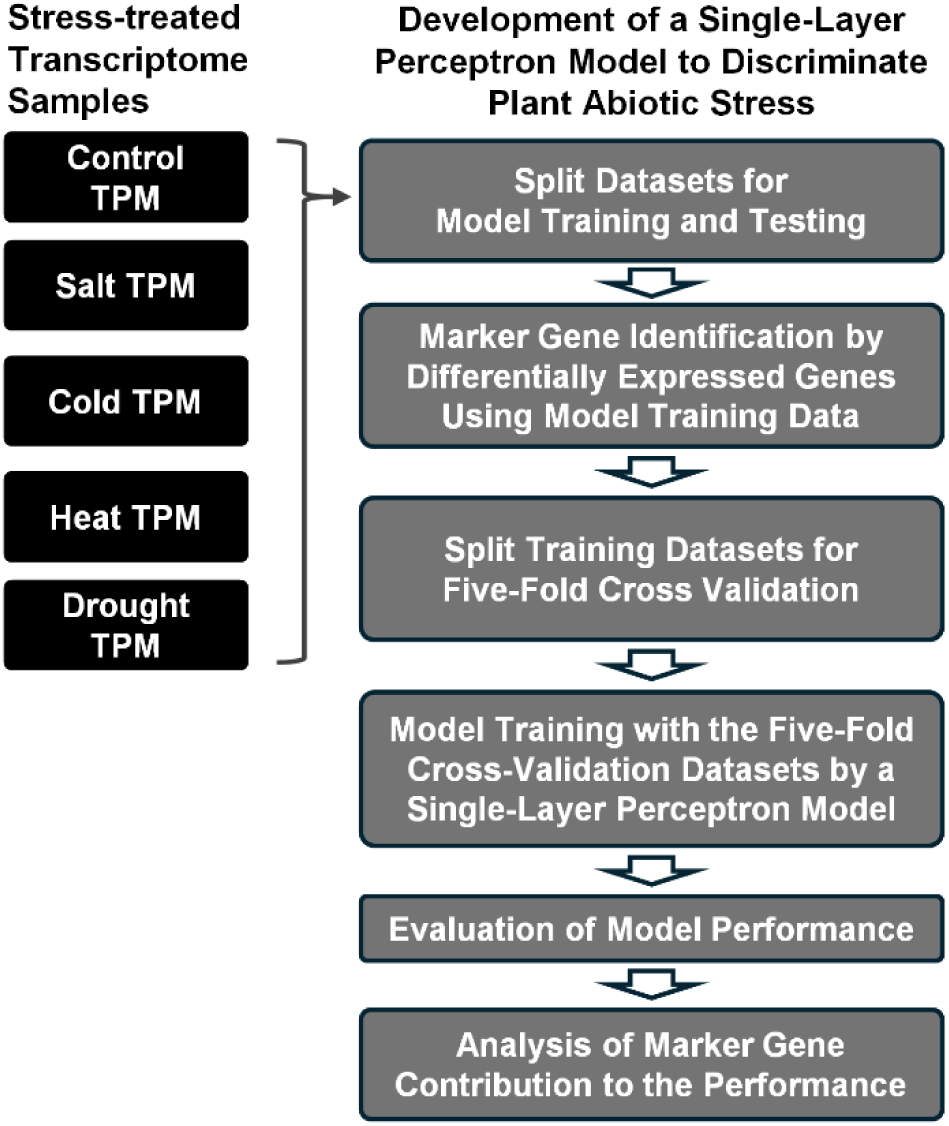
Schematic representation of the machine learning pipeline. The workflow includes data acquisition from public databases, quality trimming, transcript quantification, differential expression analysis for feature selection, and a five-fold cross-validation strategy for model training and evaluation.

### Identification of stress-specific marker genes

To identify marker genes capable of distinguishing abiotic stress responses in plants, we performed differential expression analysis using DESeq2. By comparing 120 stress-treated samples against 120 common control samples, we identified a robust set of differentially expressed genes (DEGs) with high statistical significance. Specifically, we detected 1,017 (salt), 517 (cold), 917 (heat), and 1,703 (drought) upregulated DEGs, alongside 445, 954, 1,006, and 2,587 downregulated DEGs, respectively (Figure 2A). To isolate stress-specific marker genes, we utilized Venn diagram analysis (Figure 2B), which revealed 581 (salt), 287 (cold), 632 (heat), and 1,076 (drought) unique upregulated DEGs. Similarly, unique downregulated DEGs were identified as 81 (salt), 319 (cold), 320 (heat), and 1,626 (drought). These unique DEGs were subsequently prioritized as candidate markers for stress classification.

**Figure 2.**
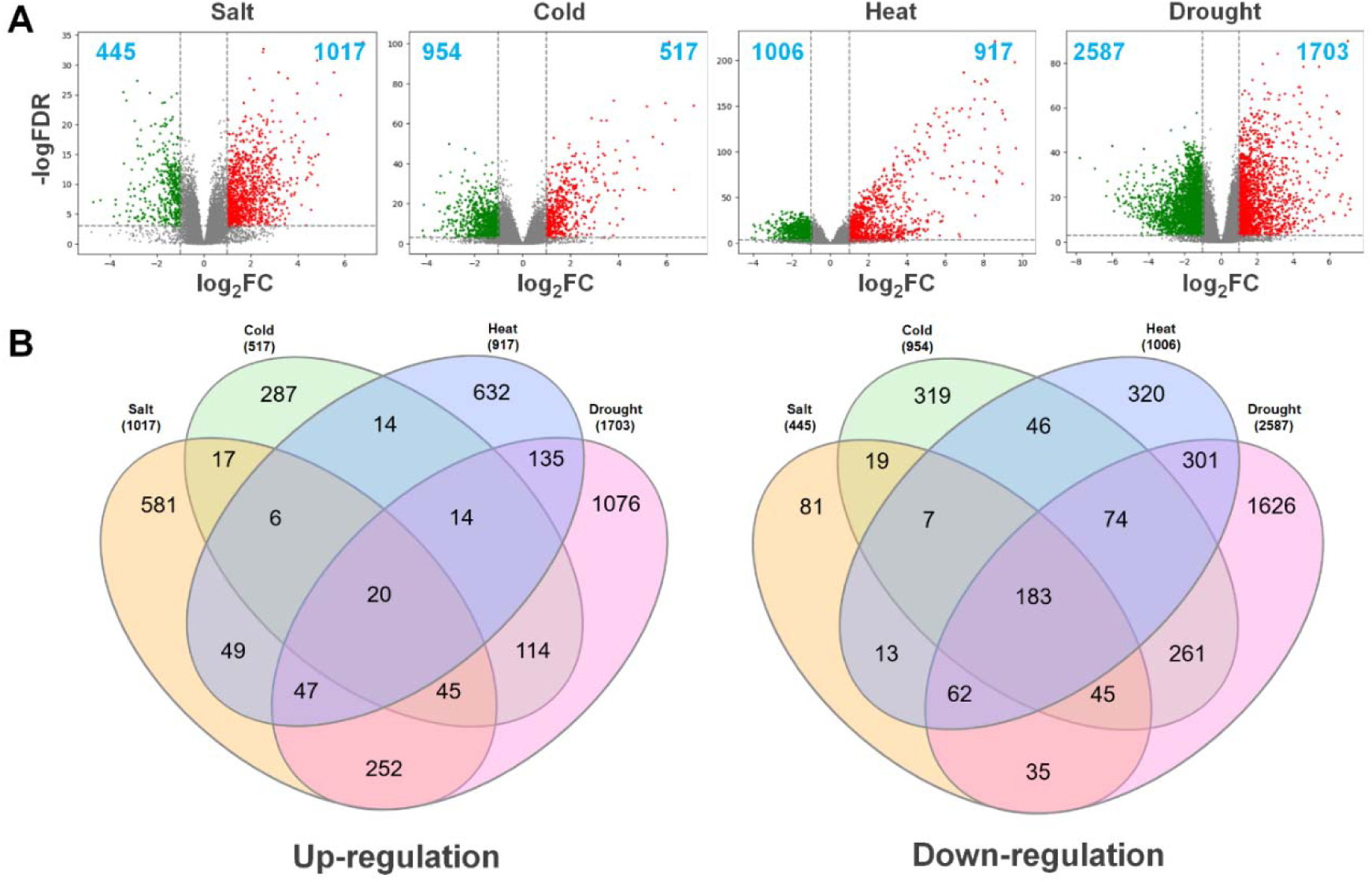
Identification of stress-responsive differentially expressed genes (DEGs). (A) Volcano plots of up-regulated and down-regulated DEGs identified for each abiotic stress condition (salt, cold, heat, and drought) compared to controls. (B) Venn diagrams showing the overlap of DEGs across the four stress conditions, highlighting the unique stress-specific DEGs used for marker gene selection.

Gene Ontology (GO) enrichment analysis of stress-specific DEGs for biological processes showed that ‘Response to stress’ was the most abundant category across all four stress conditions (Figure 3). Similarly, ‘Response to chemical’ and ‘Cellular response to stimulus’ were consistently highly ranked. When analyzing specific stress types, unique enrichment patterns were identified. ‘Cellular response to chemical stimulus’ showed the highest fold enrichment for salt stress, while ‘Response to abiotic stimulus’ was dominant for cold stress. Notably, ‘Heat acclimation’ and ‘Translation’ exhibited the greatest fold enrichment for heat and drought stress, respectively. These results confirm that identifying marker genes from large-scale datasets using DEGs effectively captures biologically relevant stress responses, supporting their utility as diagnostic markers.

**Figure 3.**
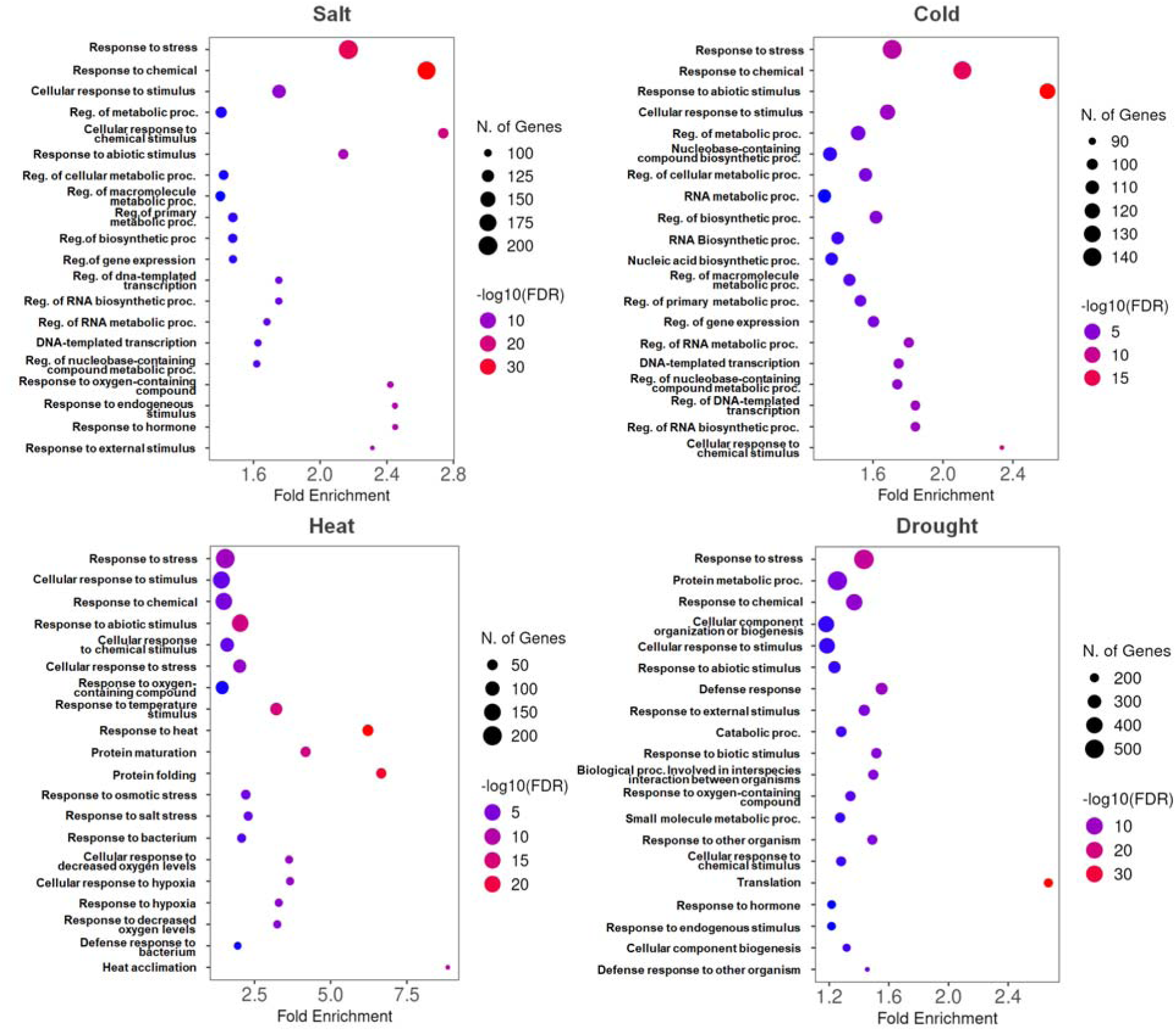
Gene Ontology (GO) enrichment analysis of stress-responsive genes. Top enriched Biological Process GO terms for the DEGs identified under salt, cold, heat, and drought stress. Significance was determined using a False Discovery Rate (FDR) threshold of 0.05.

### Dimensionality reduction and feature selection

Although initial DEG filtering reduced the feature space, the remaining 4,922 stress-specific DEGs represented an impractically large set for a diagnostic panel. To construct a concise and balanced marker set, we randomly down-sampled the pool to 40 up-regulated and 40 down-regulated genes per stress type, yielding a final set of 320 markers (Figure 4A). To evaluate the discriminatory power of this subset, we compared clustering performance using PCA and t-SNE across three input levels, all genes, total DEGs, and the 320 selected markers (Figure 4B). PCA failed to distinctly separate stress conditions. While t-SNE improved clustering, significant overlap persisted. These results indicated that unsupervised dimensionality reduction was insufficient for precise classification, requiring a supervised machine learning approach.

**Figure 4.**
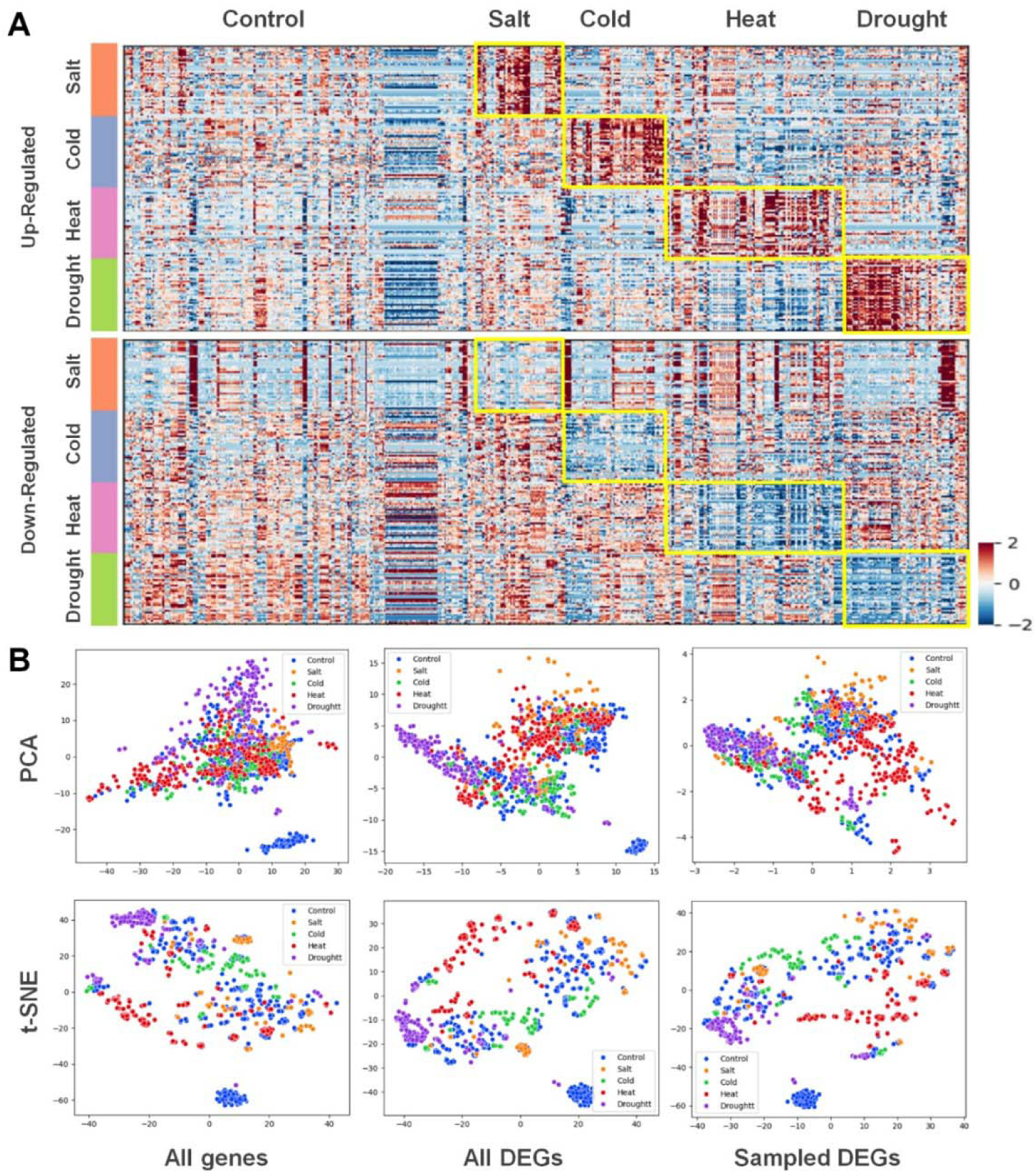
Feature selection and dimensionality reduction of transcriptomic data. (A) Heatmap visualization of the 320 selected marker genes (40 upregulated and 40 downregulated genes per stressor) across all curated samples. (B) Comparison of sample clustering using Principal Component Analysis (PCA) and t-distributed Stochastic Neighbor Embedding (t-SNE) based on all detected genes, all non-redundant DEGs, and the final 320 marker genes.

### Model architecture and performance evaluation

To develop a machine learning model, we tested a multilayer perceptron (MLP) with varying numbers of hidden layers. We tested various hyperparameter combinations of the MLP and identified that a single fully connected hidden layer (i.e., a one-hidden-layer MLP, hereafter referred to as a single-layer perceptron) yielded optimal performance (Figure 5A). To accommodate potential multi-stress conditions, the architecture used sigmoid activation at the output layer to enable multi-label classification. However, as model development was restricted to single-stress-treated samples, the class with the highest sigmoid probability was assigned as the predicted label.

**Figure 5.**
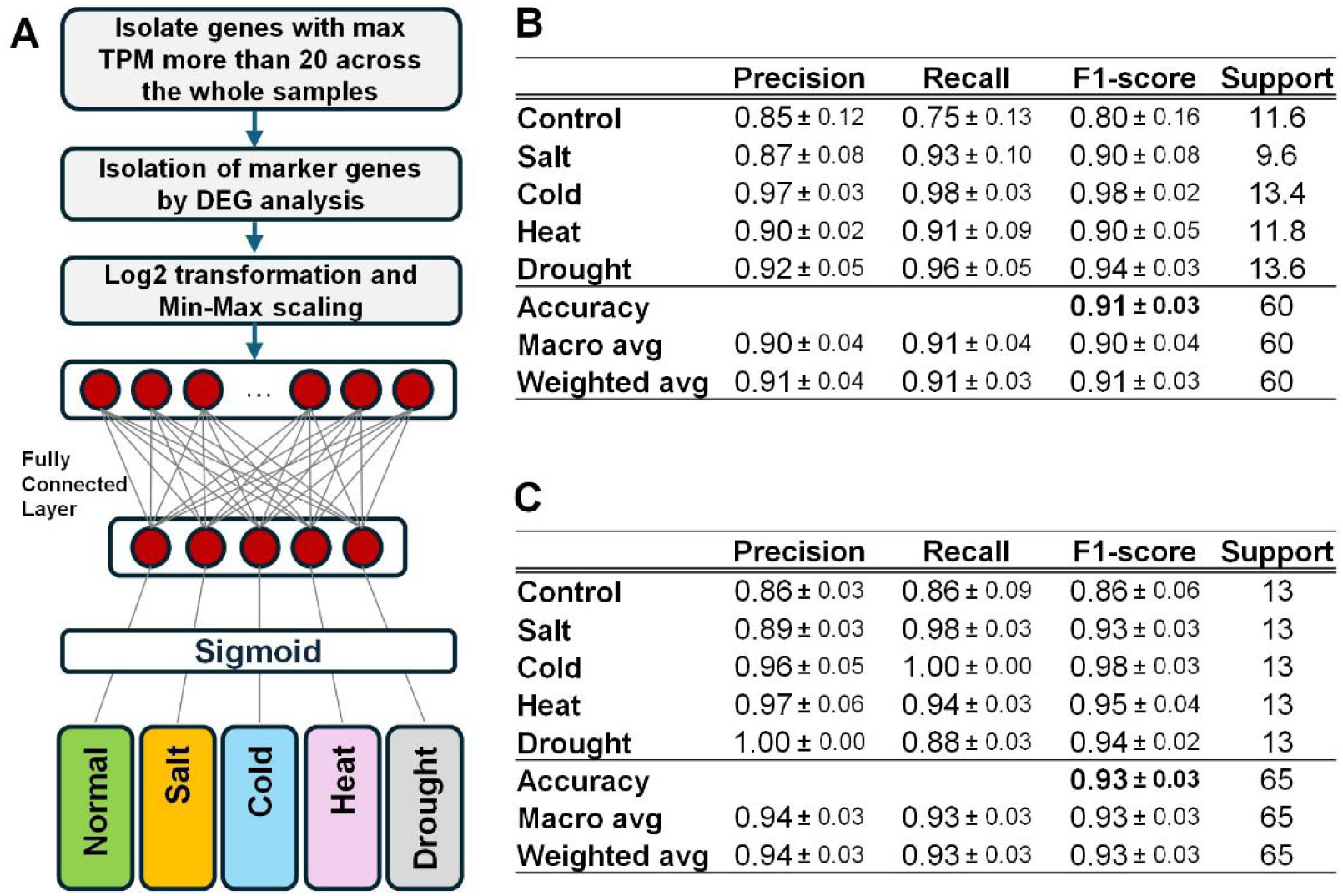
Model architecture and classification performance. (A) Architecture of the single-layer perceptron model used for stress classification. (B) Confusion matrix and performance metrics (precision, recall, and F1-score) obtained from 5-fold cross-validation on the training set. (C) Performance evaluation on the independent test set of 65 samples, demonstrating high accuracy and model generalizability.

Five-fold cross-validation demonstrated robust performance (Figure 5B), with a macro-average F1-score of 0.90 ± 0.04 and an overall accuracy of 0.91 ± 0.03. Among the classes, Cold stress achieved the highest F1-score (0.98 ± 0.02), reflecting distinct transcriptomic signatures. Conversely, Control samples exhibited the lowest F1-score (0.80 ± 0.16). The discrepancy between high precision (0.85 ± 0.12) and lower recall (0.75 ± 0.13) suggests occasional misclassification of control samples as stressed. This may reflect the inherent heterogeneity of control transcriptomes, which were pooled from experiments conducted under different baseline conditions across multiple studies. To rule out data leakage from DEG selection, we evaluated the model on 65 independent test samples that were excluded from the feature selection process (Figure 5C). If there were data leakage, the test results from the independent test samples were expected to be significantly lower than those from five-fold cross-validation. However, the resulting accuracy and F1-score were 0.93, confirming the model’s generalizability.

### Validation of the marker gene selection method and analysis of key marker genes

To determine the optimal number of features and the validity of random selection, we evaluated model performance across varying feature set sizes using 300 iterations of randomly selected marker genes. We tested total set sizes of 40, 80, 160, and 320 genes, composed of 5, 10, 20, and 40 up- and down-regulated DEGs per stress condition, respectively (Figure 6A). Average accuracy in five-fold cross-validation improved from 0.83 (40 genes) to 0.87 (80 genes) and 0.90 (160 genes), saturating at 0.91 with 320 genes. This confirms that the 320-gene set used in our final model is sufficient for optimal performance.

**Figure 6.**
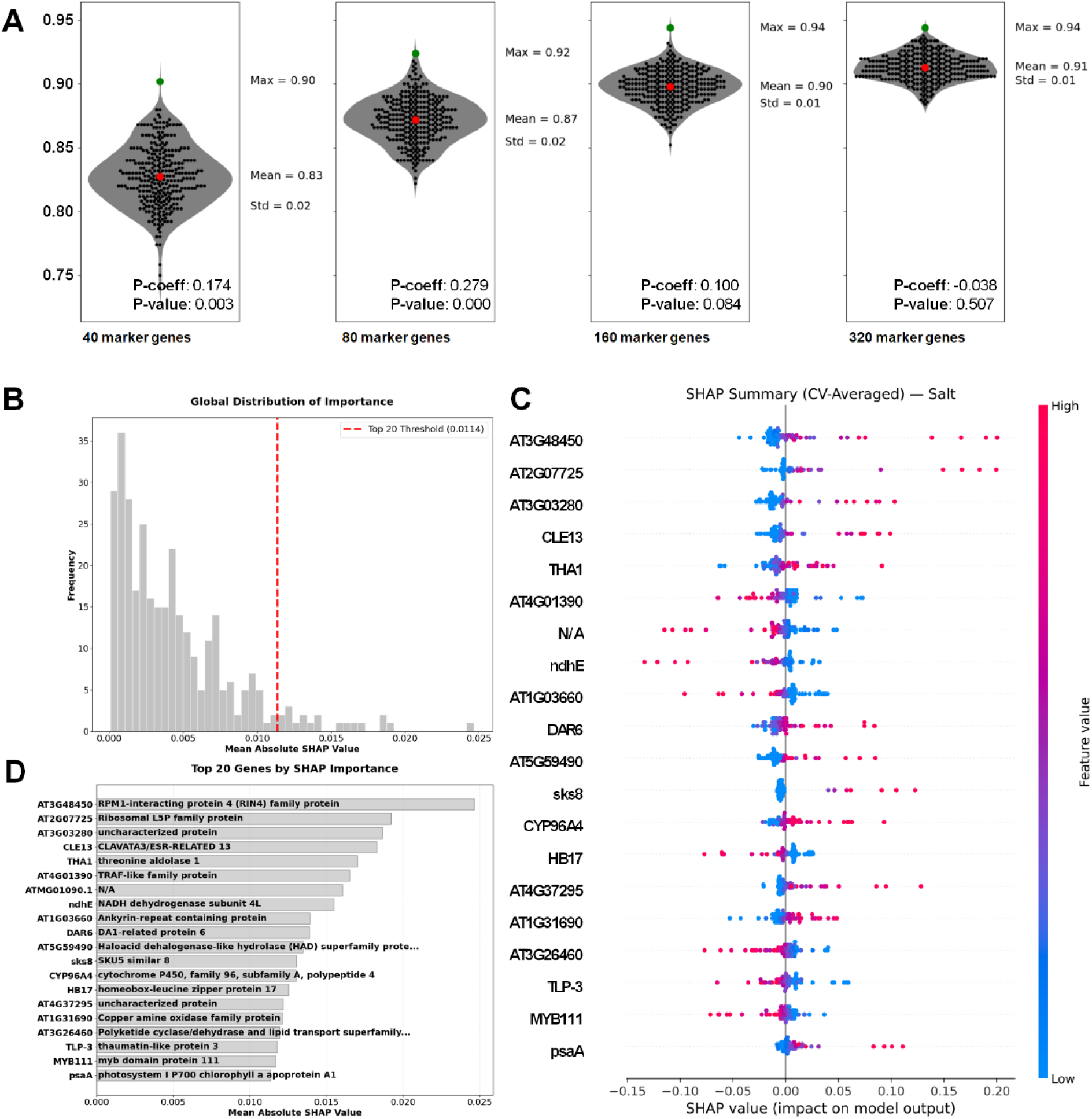
Assessment of model performance stability across varying marker gene set sizes and marker gene importance with SHAP analysis. (A) Accuracy distributions for 300 iterations of random feature selection are shown for 40, 80, 160, and 320 marker genes. Dots represent individual 5-fold cross-validation results. Summary statistics include maximum (green), mean (red), and standard deviation. Pearson correlation coefficients (P-coeff) and P-values indicating the concordance between cross-validation and independent test-set accuracy are provided for each condition. (B) Distribution of mean absolute SHAP values. (C) SHAP values of the top 20 marker genes. (D) Mean absolute SHAP values and function of the top 20 marker genes.

We further assessed whether selecting the specific gene subsets that yielded the highest cross-validation accuracy would outperform random selection. While the “best” subsets achieved accuracies of up to 0.94, a Pearson correlation test between cross-validation scores and performance on 65 independent test samples revealed a weak correlation (r < 0.279) across all set sizes. This lack of correlation indicates that optimizing for specific gene subsets in cross-validation does not guarantee generalizability to independent data, thereby supporting the robustness of our random selection approach.

To evaluate the contribution of individual marker genes to the classification of the four plant abiotic stresses and one control, we performed a global feature importance analysis using SHAP. Most marker genes had SHAP values that were less than half that of the top-performing gene (Figure 6B). The top 20 genes with the highest mean absolute SHAP values were identified, and their functions were investigated (Figure 6C and 6D, Supplementary Figures S1-4). The most important gene to discriminate salt stress was the RPM1-interacting protein 4 (RIN4) family protein (Figure 6D). Similarly, UDP-Glycosyltransferase superfamily protein and xyloglucan endotransglucosylase/hydrolase 13 were the most important genes for discriminating between cold and heat stress, respectively (Supplementary Figures S1 and S2). In drought and control conditions, the most important gene was lipid transfer protein 4, indicating that the gene’s up- and down-regulation can distinguish the two conditions (Supplementary Figures S3 and S4).

### Evaluation of Multi-Stress Samples

To assess model performance under complex conditions, we evaluated samples subjected to combined stress treatments. We analyzed eight samples treated with combined Salt and Heat stress (36–38) and three treated with Heat and Drought stress (39). The model identified both stress signatures in the Salt+Heat samples. However, in the Heat+Drought samples, only the drought signature was detected (Figure 7). As detailed in the Discussion section, this partial detection is likely due to the specific intensity thresholds used in the heat treatment in that dataset. Although formal statistical validation was precluded by the very small sample sizes (n = 8 and n = 3, respectively), the dual classification observed in the Salt+Heat group provides preliminary evidence that models trained on single-stress data may retain the ability to generalize to multi-stress conditions when stress intensities are sufficient.

**Figure 7.**
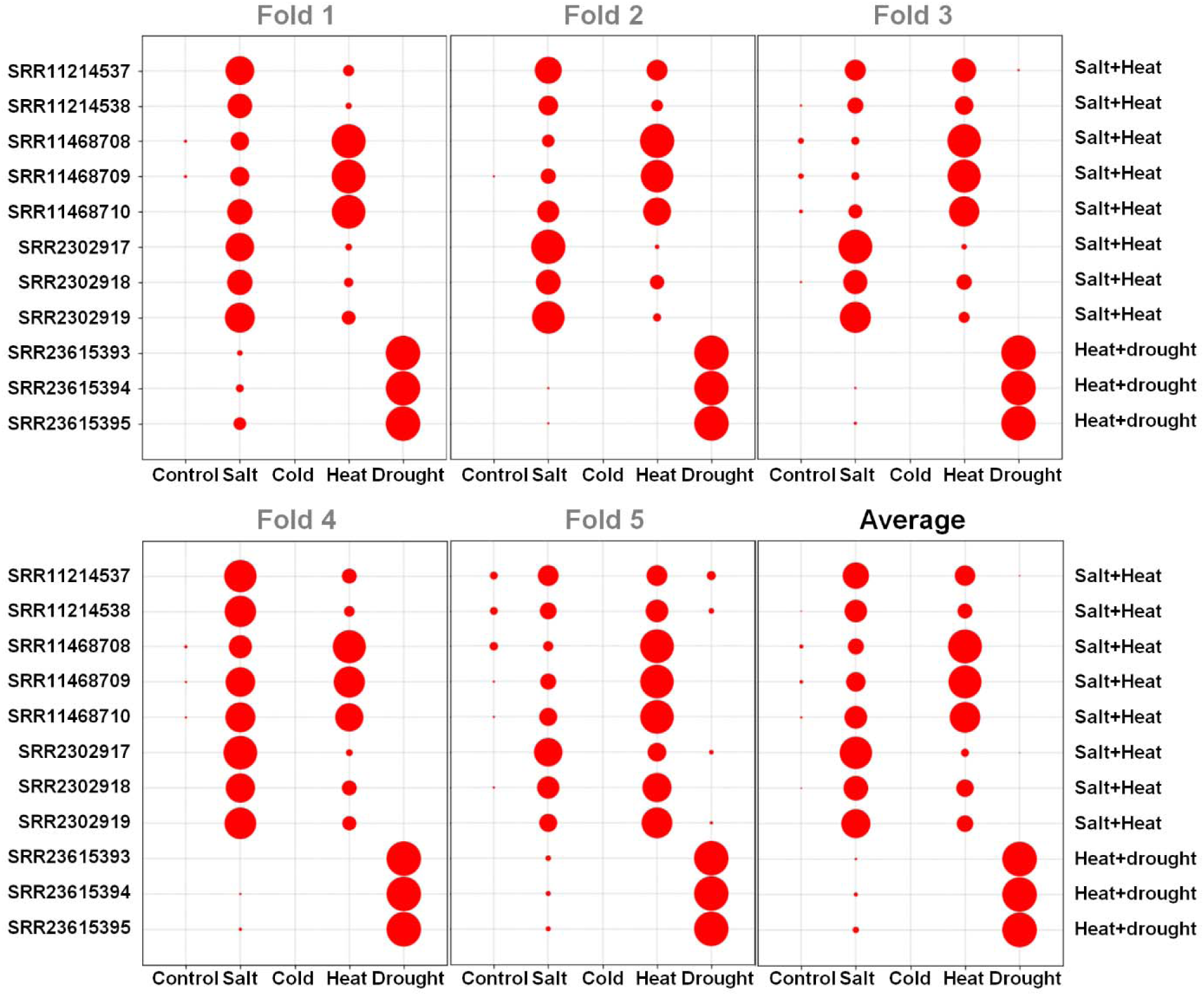
Model performance on multi-stress combinatorial samples. Visualization of sigmoid output probabilities for samples subjected to simultaneous stressors (e.g., salt + heat and heat + drought). Circle size represents the magnitude of the predicted probability for each stress class. The model successfully identified both stressors in salt-heat combinations but prioritized drought signatures in heat-drought samples.

## Discussion

Our results demonstrate that plant abiotic stress can be accurately classified using a machine learning model trained on transcriptomic profiling data. To our knowledge, this is the first study to report a method capable of distinguishing among multiple types of abiotic stress in plants. Transcriptomic data provide a powerful resource for early detection, as changes in gene expression occur well before visible physiological symptoms appear (40,41). For example, Kawasaki et al. (2001) reported detectable transcriptomic alterations as early as 15 min after salt exposure (21). These findings support the utility of transcriptome-based approaches for the sensitive and early identification of plant stress responses.

While abiotic stress is a major constraint on crop productivity (42), existing detection methods, such as thermal infrared and visible-spectrum imaging, have limitations. Visible imaging detects stress only after phenotypic symptoms emerge, and while thermal imaging can identify early physiological changes, neither approach can definitively identify the specific underlying cause (43,44). Our method addresses this limitation by enabling early detection while simultaneously pinpointing the specific stressor, offering a more actionable strategy to mitigate yield loss.

We used *Arabidopsis* because extensive RNA-seq datasets representing diverse stress treatments are publicly available. These datasets, however, were generated for heterogeneous purposes and were not optimized for stress classification. To curate clear training examples, we excluded samples exposed to confounding stressors, such as pathogens or herbicides, and limited the dataset to leaf-containing samples. This allowed us to assemble a set of single-stress, leaf-derived transcriptomes for model training. We did not filter samples by species, ecotype, or mutant background, assuming that stress-responsive transcriptional patterns would be broadly conserved. Any gene expression differences attributable to genotype variation would be assigned low weights during model training and thus minimally influence classification performance. One notable limitation of the current dataset is the heterogeneity of control samples, which were pooled from multiple independent experiments conducted under varying baseline conditions. This likely contributes to the comparatively lower F1-score observed for the control class, as the model must generalize across a wide range of “unstressed” transcriptional states. Future datasets with standardized control conditions would be expected to improve classification performance for this class.

A key challenge in realistic agricultural settings is the occurrence of combined stresses. Our model successfully detected both stressors in samples treated with Salt and Heat. However, in samples treated with combined Heat and Drought, the model detected only the drought signal. We attribute this discrepancy to inconsistencies in the definition of “stress” across public datasets. An analysis of the experimental metadata reveals that the Heat–Salt samples were treated at clear stress-inducing temperatures: 35°C (SRR11214537–38) (36), 33°C (SRR11468708–10) (37), or 43°C (SRR2302917–19) (38). In contrast, the Heat–Drought samples (SRR23615393–95) were exposed to only 27°C (39). As the model was trained on samples typically treated at ≥ 33°C, it likely classified the 27°C condition as non-stress (normal) relative to the heat signature. This observation highlights that multi-stress classification requires training data with stress thresholds that align with the specific definitions of stress in the target environment.

Although transcriptomic profiling provides high-resolution information about plant stress responses, it remains costly and time-consuming, limiting its use in cases requiring immediate decision-making. Nevertheless, we propose two practical applications. First, transcriptome-based stress classification can serve as a high-confidence labeling strategy for training other machine learning models that rely on cultivation metadata or imaging data. As machine learning approaches for plant stress detection continue to advance (19,45,46), the accuracy of training labels remains a critical determinant of model performance. Our method provides evidence-based, biologically grounded labels that can enhance downstream model reliability. Second, transcriptome-based classification can support precision breeding by distinguishing stress-resistant from stress-tolerant genotypes. As climate change intensifies, stress-tolerant cultivars may maintain survival but suffer yield penalties, making it essential to identify truly stress-resistant lines. Our approach provides a quantitative framework for phenotyping breeding populations using molecular stress signatures.

In summary, this study presents the first machine learning model capable of classifying multiple abiotic stress types in plants, achieving over 91% accuracy and demonstrating the potential to identify combined stress conditions. To accommodate the broad application of the methods, we also developed an end-to-end pipeline to train models in various plant species. Future work should focus on generating high-quality training datasets from crop species and developing stress-induction protocols that reflect realistic agricultural environments. Defining crop-specific stress thresholds and designing experiments optimized for stress classification will further enhance model performance. Ultimately, this approach provides a foundation for AI-driven decision-support systems in crop management and precision breeding, offering a timely tool for addressing the challenges posed by global climate change.

## Acknowledgements

We thank all researchers who have deposited transcriptome data of abiotic stress-treated *Arabidopsis* plants in the public database.

## Supplementary data

Supplementary Figure S1. SHAP analysis of marker genes for cold stress.

Supplementary Figure S2. SHAP analysis of marker genes for heat stress.

Supplementary Figure S3. SHAP analysis of marker genes for drought stress.

Supplementary Figure S4. SHAP analysis of marker genes for control.

