## Supplementary Figure S1-S4 for "AbiOmics: An End-to-End Pipeline to Train Machine Learning Models for Discrimination of Plant Abiotic Stresses Using Transcriptomic Profiling Data"

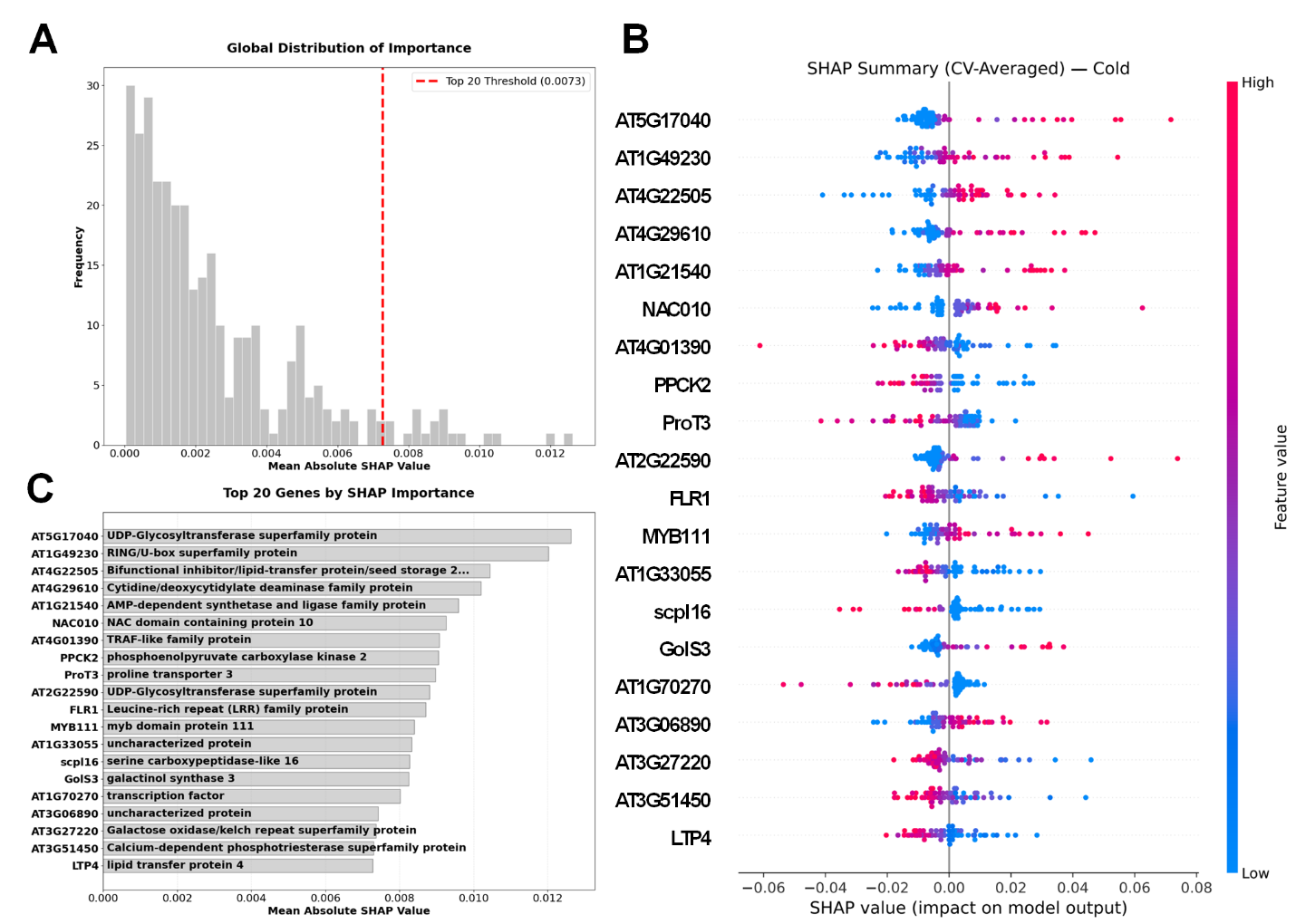

**Supplementary Figure S1. SHAP analysis of marker genes for cold stress.** (B) Distribution of mean absolute SHAP values. (C) SHAP values of the top 20 marker genes. (D) Mean absolute SHAP values and function of the top 20 marker genes.

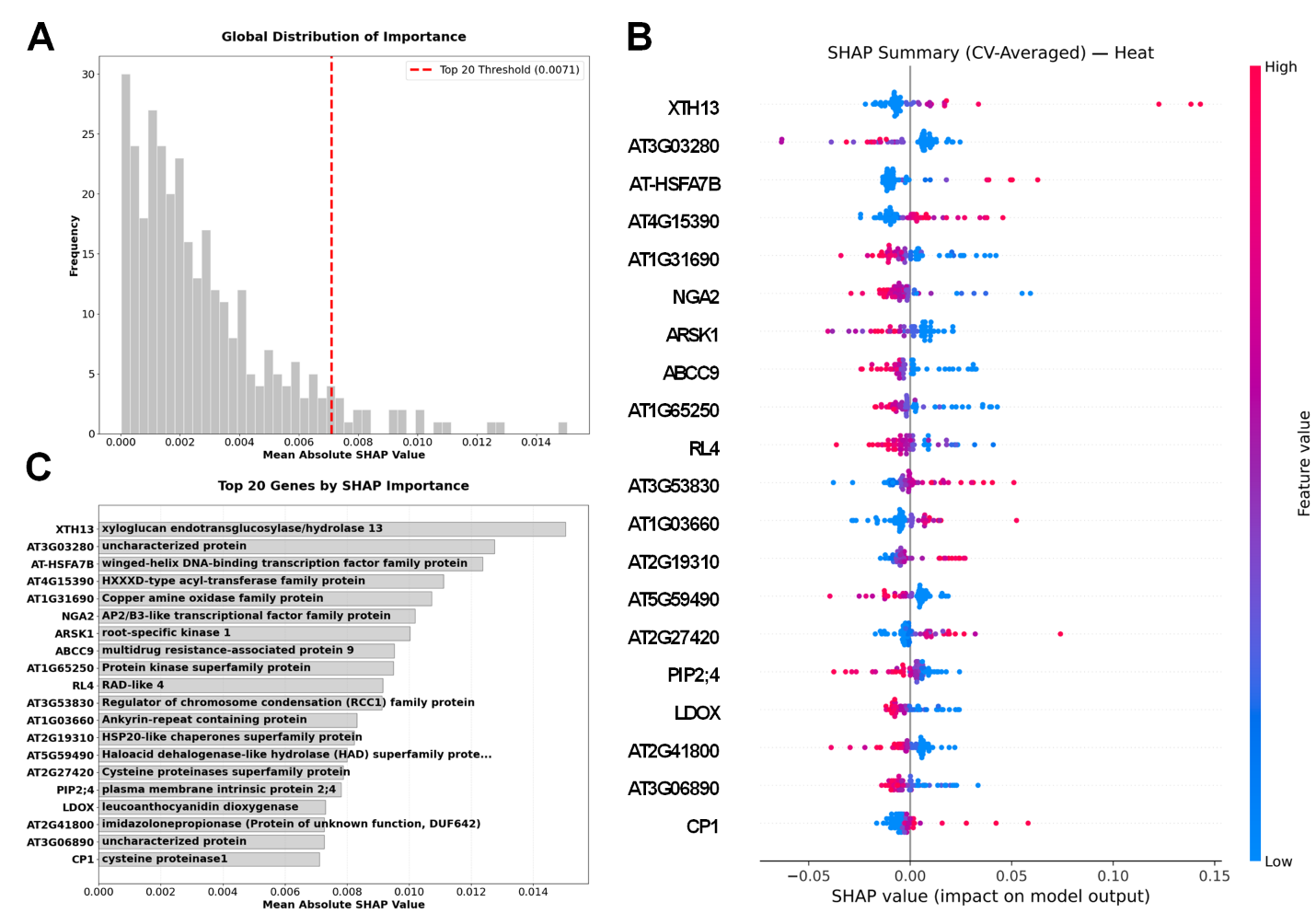

**Supplementary Figure S2. SHAP analysis of marker genes for heat stress.** (B) Distribution of mean absolute SHAP values. (C) SHAP values of the top 20 marker genes. (D) Mean absolute SHAP values and function of the top 20 marker genes.

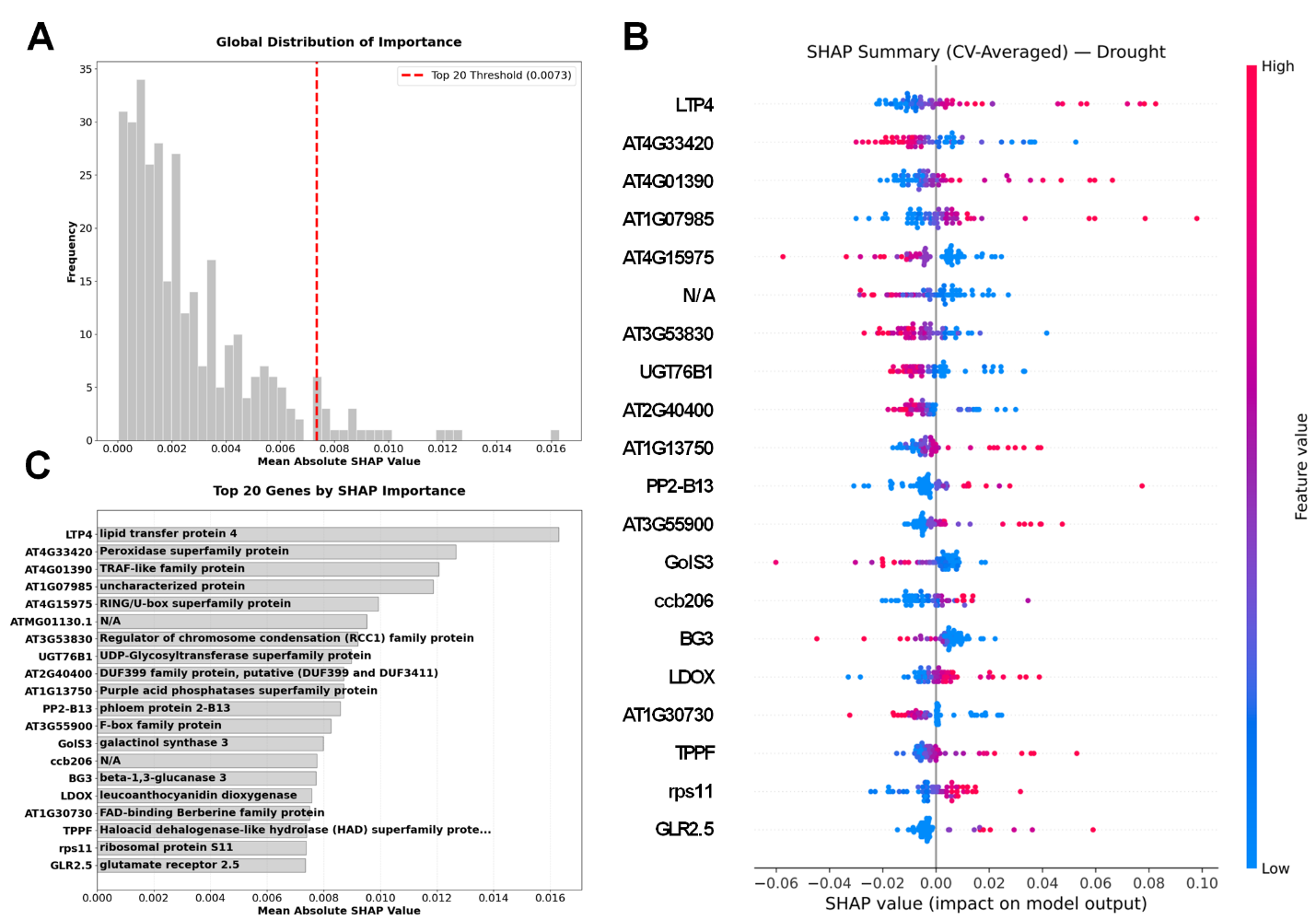

**Supplementary Figure S3. SHAP analysis of marker genes for drought stress.** (B) Distribution of mean absolute SHAP values. (C) SHAP values of the top 20 marker genes. (D) Mean absolute SHAP values and function of the top 20 marker genes.

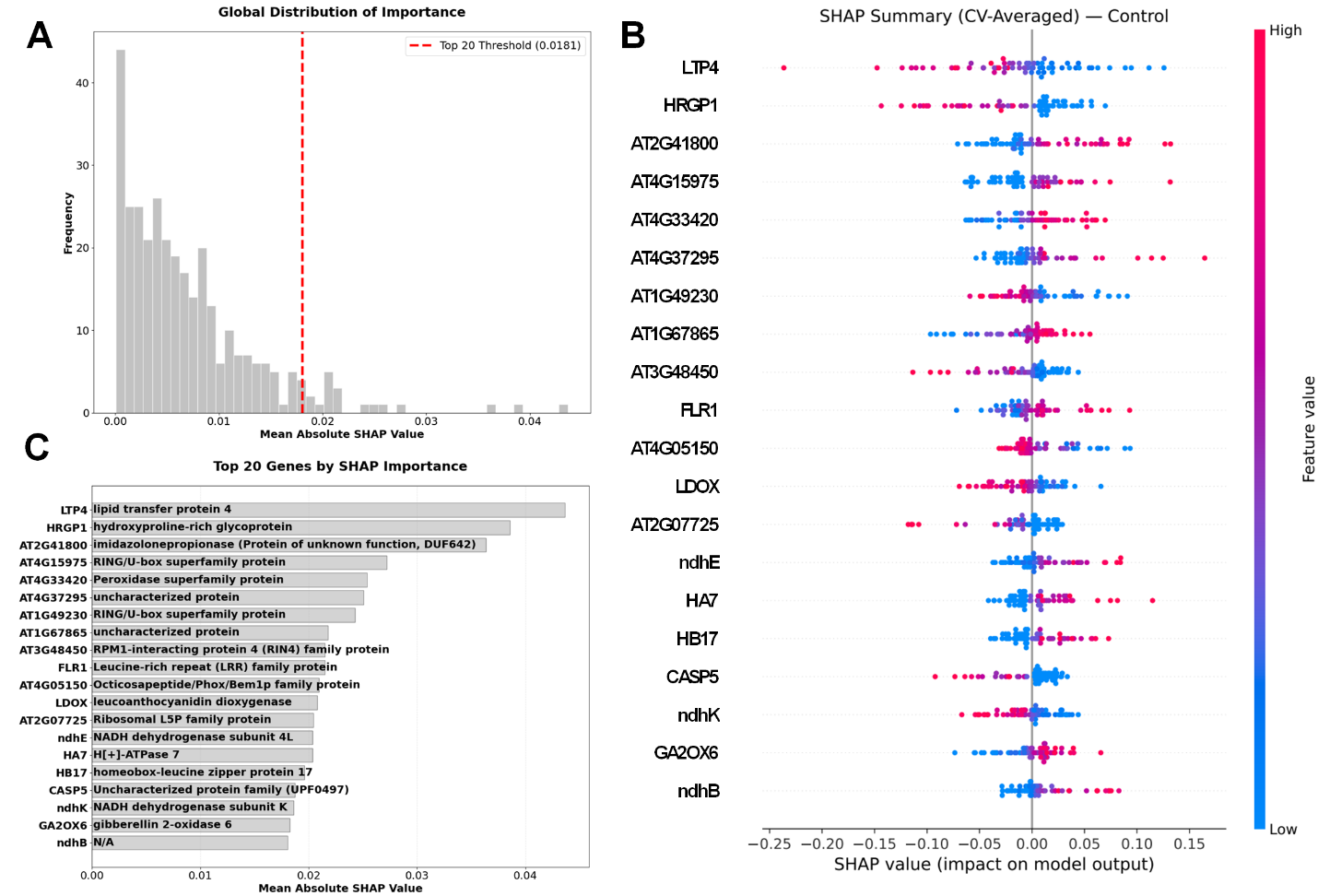

**Supplementary Figure S4. SHAP analysis of marker genes for control.** (B) Distribution of mean absolute SHAP values. (C) SHAP values of the top 20 marker genes. (D) Mean absolute SHAP values and function of the top 20 marker genes.
